# Extending long-range phasing and haplotype library imputation algorithms to very large and heterogeneous datasets

**DOI:** 10.1101/477398

**Authors:** Daniel Money, David Wilson, Janez Jenko, Gregor Gorjanc, John M. Hickey

## Abstract

**Background:** This paper describes the latest improvements to the long-range phasing and haplotype library imputation algorithms that enable them to successfully phase both datasets with one million individuals and datasets genotyped using different sets of single nucleotide polymorphisms (SNPs). Previous publicly available implementations of long-range phasing could not phase large datasets due to the computational cost of defining surrogate parents by exhaustive all-against-all searches. Further, both long-range phasing and haplotype library imputation were not designed to deal with large amounts of missing data, which is inherent when using multiple SNP arrays.

**Methods:** Here, we developed methods which avoid the need for all-against-all searches by performing long-range phasing on subsets of individuals and then combing results. We also extended long-range phasing and haplotype library imputation algorithms to enable them to use different sets of markers, including missing values, when determining surrogate parents and identifying haplotypes. We implemented and tested these extensions in an updated version of our phasing software AlphaPhase.

**Results:** A simulated dataset with one million individuals genotyped with the same set of 6,711 SNP for a single chromosome took two days to phase. A larger dataset with one million individuals genotyped with 49,579 SNP for a single chromosome took 14 days to phase. The percentage of correctly phased alleles at heterozygous loci was respectively 90.5% and 90.0% for the two datasets, which is comparable to the accuracy achieved with previous versions of AlphaPhase on smaller datasets.

The phasing accuracy for datasets with different sets of markers was generally lower than that for datasets with one set of markers. For a simulated dataset with three sets of markers 2.8% of alleles at heterozygous positions were phased incorrectly whereas the equivalent figure with one set of markers was 0.6%.

**Conclusions:** The improved long-range phasing and haplotype library imputation algorithms enable AlphaPhase to quickly and accurately phase very large and heterogeneous datasets. This will enable more powerful breeding and genetics research and application.

## Background

Here we describe the latest improvements to the Long-Range Phasing (LRP) and Haplotype Library Imputation (HLI) algorithms to phase genotypes for hundreds of thousands of individuals that have been genotyped on different platforms. Phasing genotypes is the process of inferring the parental origin of individual’s alleles. This process resolves the inheritance of chromosome segments in a population and is as such a cornerstone technique in genetics. For example, it is useful for making genotype calls, imputing genotypes, detecting phenotype-genotype associations in the presence of effects such as allele-specific expression, and inferring recombination points and demographic history [1].

The size of genomic datasets has grown rapidly in recent years as genotype data is collected on an increasing number of individuals. In agriculture this growth has been driven by the increased value of genomic selection [2–4], whereas in human genetics it has been driven by the increased power of genome-wide association studies [5–7] and genomic prediction in human medicine [8]. Examples of such large datasets include the UK Biobank [9], which has recently released genotype data from approximately half a million people [10], and the US Dairy Cattle and Irish Cattle Breeding Federation Databases that each host well over a million of genotyped animals [4,11,12].

In many cases these datasets have been collected over several years and have been genotyped using different single nucleotide polymorphism (**SNP**) arrays [4,12]. Methods such as SNPchiMp [13] have been developed to allow the manipulation of different sets of markers from multiple SNP arrays, but their main aim is to ensure that the different sets are combined correctly rather than to perform analyses of the combined dataset.

Several methods for phasing genotype data have been developed based on probabilistic methods, such as those implemented in fastPHASE [14] and Beagle [15]. Others, such as AlphaPhase [16] and findHap [17], are based on heuristic methods. Recent developments in probabilistic methods e.g., SHAPEIT3 [18] and Beagle [15], have enabled phasing of very large datasets, potentially containing over one million individuals [19]. Heuristic methods are fast when compared to statistical methods and, in many cases, more accurate. Thus, enabling their use on large datasets will be beneficial for large scale genomic studies.

AlphaPhase [16] is a heuristic method that combines LRP [20] and HLI. LRP infers parental origin of alleles by finding surrogate parents of an individual, that is individuals who likely have the same haplotype as the individual. If a surrogate parent is homozygous then it can be used to phase the individual’s genotype. When a homozygous surrogate parent cannot be found, surrogate parents of the heterozygous surrogate parent can be used. This process is repeated, with increasingly remote surrogate parents, until the individual’s genotype can be phased.

HLI infers the phase of a genotype by creating a library of haplotypes that are fully phased. Partially phased haplotypes can be fully phased by matching with library haplotypes.

Existing publicly available LRP algorithms cannot efficiently phase large datasets as finding surrogate parents amongst all individuals in a population involves comparing every individual with every other individual. Both runtime and memory usage quickly become impractical with large datasets as they scale with the square of the number of individuals.

Additionally, existing publicly available algorithms for LRP and HLI could not phase heterogeneous datasets with different sets of markers as they were not designed to cope with large amounts of missing data. Combining data from multiple SNP arrays can lead to large amounts of missing data.

In this paper we introduce improvements that allow a) LRP of large datasets and b) LRP and HLI to work with missing data. These improvements enabled us to quickly and accurately phase large heterogeneous simulated datasets. We phased one million individuals, genotyped for 49,579 SNPs, in 14 days using modest computing resources and correctly phased over 90% of alleles at heterozygous loci. We were also able to phase a dataset consisting of individuals genotyped with three different arrays and correctly phased 95% of alleles at heterozygous loci. The percentage of incorrectly phased alleles at heterozygous loci was respectively only 1.0% and 2.8% for the two examples. Our results show that it is possible to quickly and accurately phase large heterogeneous datasets and our algorithm improvements will be of benefit to those conducting large scale genomic studies.

## Methods

### Previous LRP and HLI Algorithms

Both LRP and HLI operate on genome regions called cores. A core is a set of consecutive SNPs for which phasing is being attempted. For further details see Hickey *et al.* [16].

LRP infers the phase of an individual by using other individuals known to share a haplotype with the individual. Individuals sharing a haplotype are called “surrogate parents” (shortened here to “surrogates”) and are identified by finding no opposing homozygote markers at any position within a core. These surrogates are then partitioned into either paternal or maternal surrogates of the individual using pedigree information, if it is available. If pedigree information is not available, this assignment is arbitrary.

If a surrogate is homozygous at a position then it enables phasing of the individual at that position. If no homozygous surrogate is found, then it may be possible to phase the individual by using surrogates of surrogates. This process can be continued to an arbitrary depth. In practice, the consensus of several homozygous surrogates is taken to allow for error in determining surrogates or genotype data.

HLI infers phase by matching partially phased haplotypes to a library of known haplotypes. In the existing algorithm the initial library is constructed from the fully phased haplotypes found during LRP and by adding new haplotypes as they are discovered. New haplotypes are discovered when one haplotype of an individual is inferred, because this haplotype together with genotype information determines the other haplotype of the individual. This process is iterated until no new haplotypes are found.

### Extending long-range phasing to large datasets

To address the problem of scaling LRP to large datasets we modified the algorithm so that it is performed on subsets of individuals and the results from each subset are combined. By performing LRP on subsets the runtime can be vastly reduced, because search for surrogates has quadratic runtime scaling, and in the worst case involves all-against-all search for surrogates. When datasets are very large, all-against-all search for surrogates on the full dataset is too time consuming, while splitting the data into subsets limits the search time. Subsets of individuals, without replacement, are chosen randomly so that every individual is in a subset. These subsets are then merged and HLI is run on the complete dataset. We refer to this as the sub-setting method.

Preliminary analysis showed that including individuals in multiple subsets did not offer a significant improvement in accuracy, but increased runtime significantly (data not shown). Including related individuals in subsets also decreased accuracy (data not shown).

### Extending long-range phasing and haplotype library imputation to heterogeneous datasets

The LRP algorithm was modified to enable the identification of surrogates in the presence of missing data. Missing data hinders the identification of opposing homozygotes and so has the potential to wrongly identify surrogates. To alleviate this problem we introduced an additional parameter that defines the required number of shared markers by two individuals before surrogacy is tested.

The HLI algorithm required more complex modifications. In a multiple SNP array setting it is likely that most, or even all, individuals will have been genotyped with one array, so will not have a data for all array markers. Consequently, LRP cannot infer parental origin of alleles at missing markers. We developed methods that allowed partially inferred haplotypes to be included in the haplotype library and to be used to infer other haplotypes.

Allowing for partially inferred haplotypes in the haplotype library makes matching a new partially inferred haplotype to a library haplotype much more difficult. It is necessary to ensure that the two haplotypes have enough markers with non-missing information to be confident they are the same haplotype. Thus, we added a parameter to the HLI algorithm that specifies the required number of shared alleles to match two haplotypes (Figure 1a).

**Figure 1.**
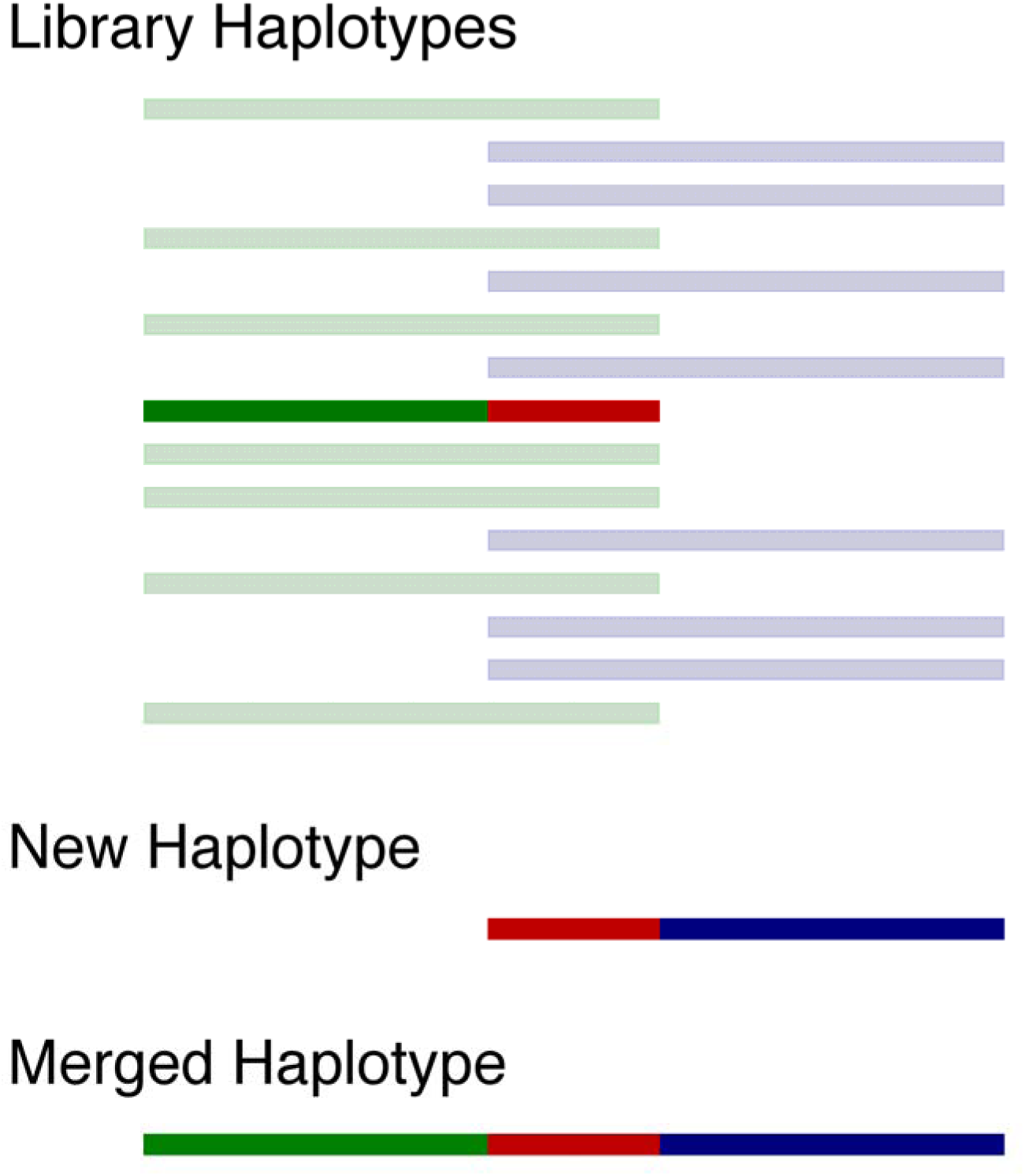
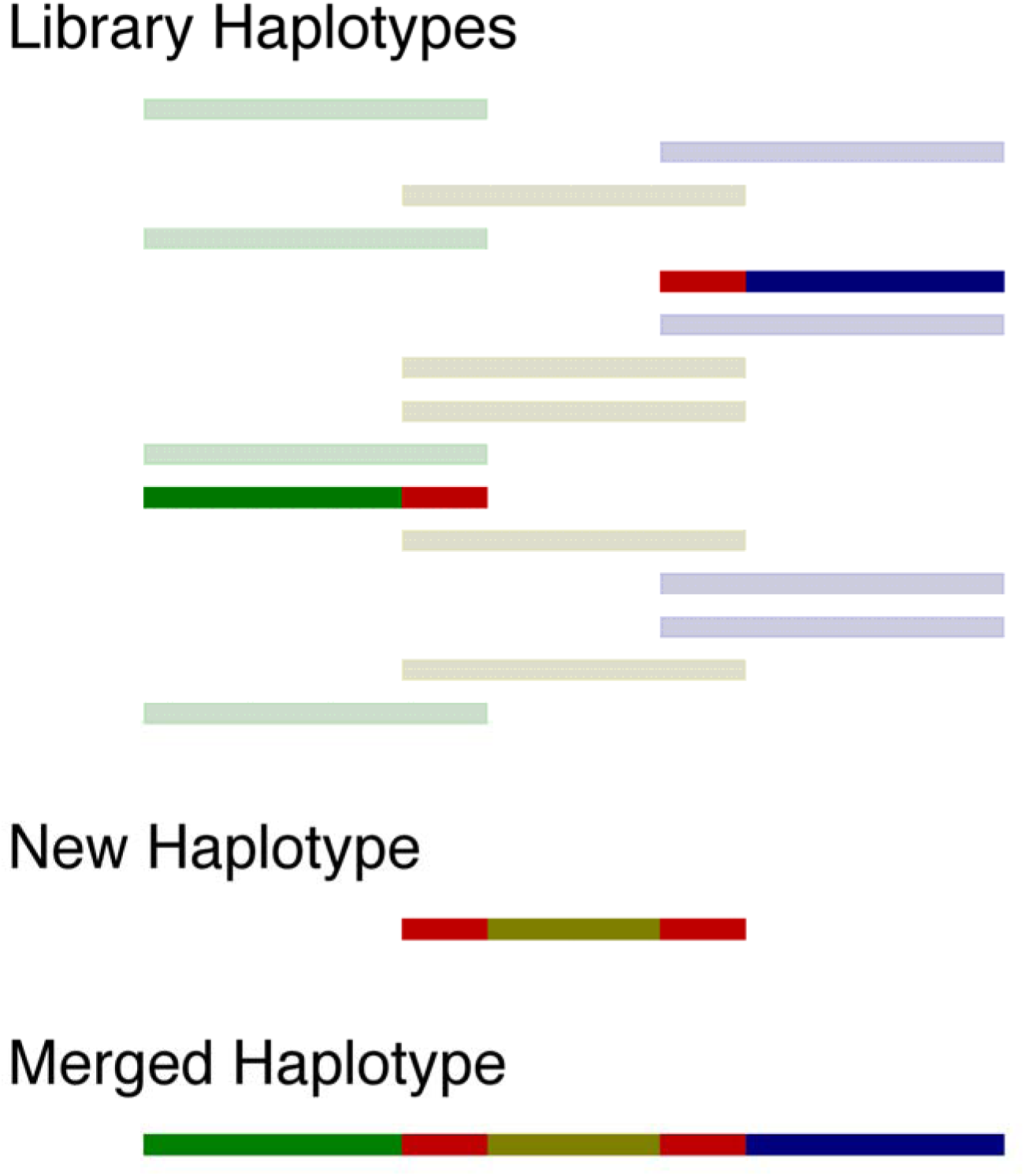
New improvements to the LRP and HLI algorithms for dealing with library haplotypes with missing data. a) In this example we have generated haplotypes using two different SNP arrays indicated by green and blue haplotypes. If the shared markers between two haplotypes are identical (shown in red) then the two haplotypes can be merged into one haplotype. To ensure the two haplotypes are the same haplotype we set a minimum number of alleles that must be shared. Note that in reality blue and green markers will both be distributed along the length of the haplotype. b) In this example we have generated haplotypes using three different SNP arrays. Finding the new purple haplotype allows us to recognise that the green, purple, and blue haplotype are actually the same haplotype.

In some cases it is possible that the new haplotype matches more than one library haplotype. In these cases it is possible that the new haplotype identifies duplicated library haplotypes (Figure 1b). In this situation we merge the new haplotype and library haplotypes, replace the incomplete library haplotypes with the merged haplotype, and update individuals known to carry the original incomplete library haplotypes.

If a new haplotype matches multiple library haplotypes and these matches cannot be the same haplotype, due to opposing homozygotes between the library haplotypes, then we add the new haplotype to the library.

### Software engineering the new algorithms

Several changes were made to AlphaPhase to optimise it for speed and memory use. AlphaPhase was modified to store haplotypes and genotypes as bits and to use bit operations to operate on multiple SNPs at once wherever possible. It was also parallelised in several places to exploit high performance computing clusters.

### Test Datasets

The performance of our new improvements to the LRP and HLI algorithms were tested on large and heterogeneous simulated datasets.

### Simulated Data

AlphaSim [21] was used to simulate test datasets. We followed the simulation scheme from [22] which we describe briefly and show in Figure 2. AlphaSim first uses MaCS [23] to simulate base population haplotypes. We simulated a single ‘breed’ that split into three breeds 400 generations ago. 50 generations ago each of these breeds split again into either three or four breeds to give ten breeds.

**Figure 2.**
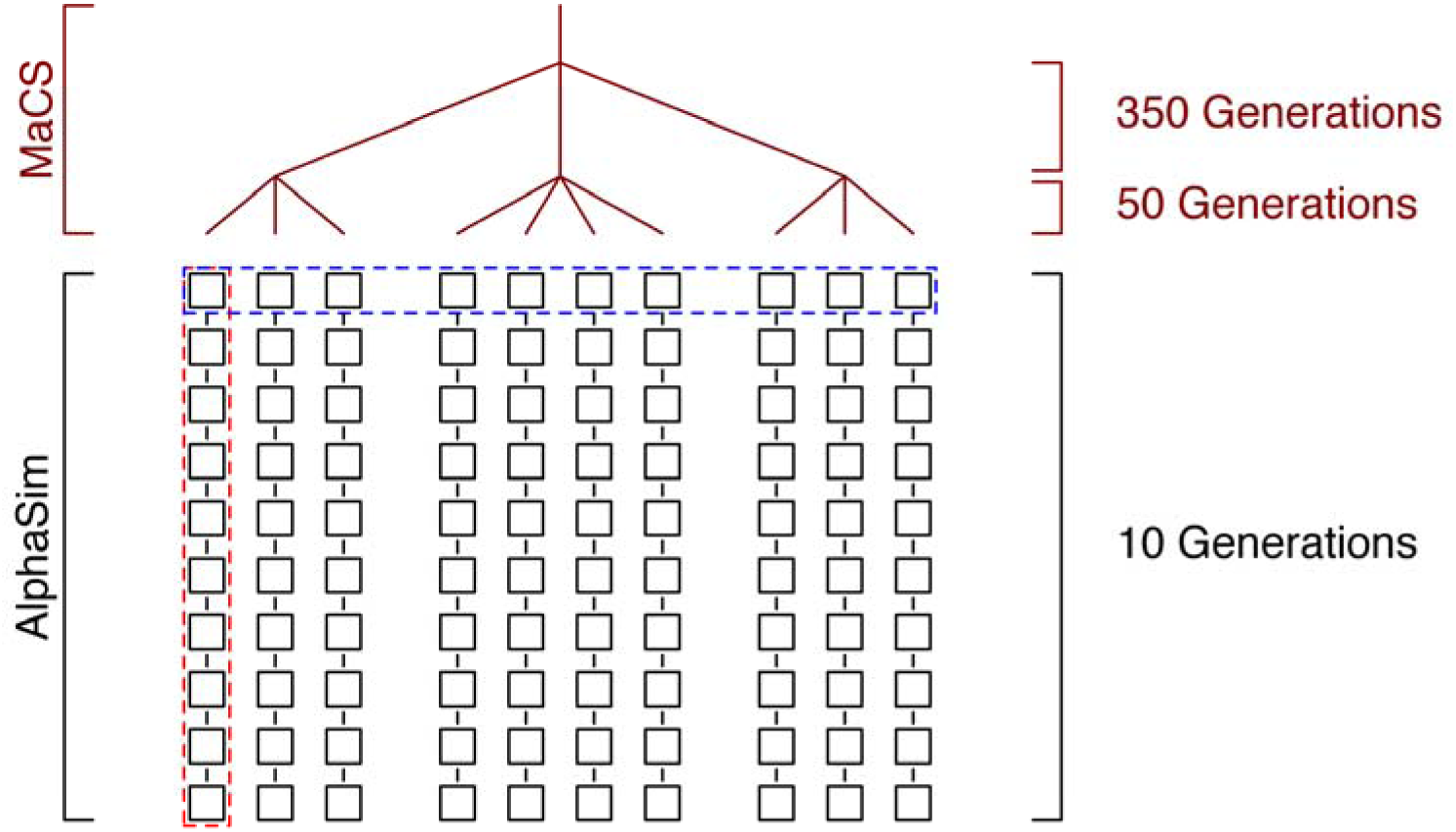
Simulation structure. MaCS is used to simulate a base population. This base population is generated from a single ‘breed’ that split into three breeds 400 generations ago. 50 generations ago each of these breeds split again into either three or four breeds to give ten breeds. Each of these ten breeds then undergoes ten generations of selection using AlphaSim. The dotted blue line shows an example for the “per generation” scenario, while the dotted red line shows an example for the “per family” scenario.

AlphaSim was then used to simulate two datasets consisting of ten equal-sized ‘breeds’ and ten generations of selective breeding were simulated for each of these ‘breeds’ (Figure 2). Selection was based on a single trait that had 10,000 quantitative trait nucleotides with normally distributed effects. For the first dataset, for each breed and for each generation, we selected 25 sires and 500 dams and generated 1,000 offspring. This resulted in a dataset of 100,000 animals (**100k dataset**). The second dataset was created using 10,000 offspring for each breed and for each generation to create a total dataset of one million animals (**one million dataset**). For both datasets one chromosome worth of SNP data was generated and SNPs with a minor allele frequency of at least 0.05 were chosen as possible candidates for inclusion on SNP arrays.

SNPchiMp [13] was used to obtain information on the SNPs on different arrays and the overlap between arrays. Across the bovine arrays there are 8,771 unique SNPs on chromosome 1. We selected this number of SNPs from the candidate SNPs generated by AlphaSim and then assigned SNPs to different arrays following the same pattern as reported by SNPchiMp for bovine arrays.

We then used the assigned arrays to create scenarios (Table 1), where individuals were genotyped with different arrays. There were two scenarios with **Homogeneous Arrays**, where all individuals were genotyped with either the medium density (**MD**) Bovine Illumina 50Kv2 or the high density (**HD**) Illumina HD SNP arrays. There were five scenarios with **Heterogeneous Arrays**, where individuals were genotyped with a set of partially overlapping combinations of SNP arrays. Three of these scenarios were based on different MD chips. The **Two Illumina** scenario included two different versions of the Illumina MD chip (Illumina 50K v1 and Illumina 50K v2). The **Two Mixed** scenario combined one Illumina chip (Illumina 50K v2) and one other chip (IDBv3). The **Three MD** scenario combined the Illumina 50Kv2 chip with the IDBv3 chip and the GSeekHD chip. The **mixed MD / HD** scenario combined a MD Illumina chip (Illumina 50K v2) with a HD Illumina chip (Illumina HD).

**Table 1.** The different genotyping scenarios tested.

| Scenario | Description |
| --- | --- |
| Illumina 50Kv2 | Illumina 50Kv2 (all) |
| Illumina HD | Illumina HD (all) |
| Two Illumina | Illumina 50Kv1 and Illumina 50Kv2 in a 1:1 ratio |
| Two Mixed | Illumina 50Kv2 and IDBv3 in a 1:1 ratio |
| Three MD | Illumina 50Kv2, GSeekHD and IDBv3 in a 1:1:1 ratio |
| Mixed MD/HD | Illumina 50Kv2 and Illumina HD in a 9:1 ratio |
| Ten Array | Five MD and Five HD with equal numbers of individuals genotyped on each array |

We created a further scenario that was not based on existing arrays, as in the future it is likely that individuals will be genotyped on a wider range of SNP arrays. The **Ten Array scenario** comprises five HD arrays and five MD arrays. We based the first HD and first MD arrays on the Bovine Illumina HD and Bovine Illumina 50Kv2 arrays, respectively.

We then created three further HD and three further MD arrays based on these two arrays by splitting SNPs into four categories, those on both the original HD and MD arrays (**HD/MD set**), those just on the HD array (**HD set**), those just on the MD array (**MD set**) and those on neither (**unused set**). We removed ten percent of the SNPs from each of the HD/MD, HD and MD sets and replaced them with randomly sampled SNPs from the unused set. From these new sets we then created a second HD and second MD array. We generated the third and fourth HD and MD arrays from the previous arrays in the same way. Rather than removing the exact number of SNPs that were to be added (or removed) we sampled based on a probability that resulted in the expected number of SNPs being added (or removed). Because of this sampling the resulting arrays were of slightly different in size.

We created the fifth HD and MD arrays to represent arrays from a different family of arrays, perhaps from a different manufacturer. These arrays were simulated in a similar way to the other arrays but by removing and replacing fifty percent of the SNPs from the first HD and MD arrays.

For all scenarios we simulated individuals as being genotyped on different arrays by assigning them to arrays in proportions we might expect to see in real datasets (Table 1).

### Phasing parameters used for AlphaPhase

AlphaPhase has several parameters that control phasing of alleles. Two of these were expected to have a significant effect on the performance of AlphaPhase – the existing parameter controlling core length (defined as the number of SNP in each core) and a new parameter that controlled the size of phasing subsets to speedup phasing of a large dataset.

The length of the cores can have a significant effect on phasing accuracy [16]. To find the best core length for both of the MD and HD scenarios we tested different core lengths. For the Illumina 50Kv2 scenarios we tested core lengths in the same range as those tested in Hickey *et al.* [16] for a similar size array: 50, 100, 200, 500, and 1,000 SNPs. For the Illumina HD scenario we tested core lengths of 500, 1,000, 2,000, 5,000, and 10,000 SNPs because the Illumina HD array contains approximately ten times as many SNPs as the Illumina50Kv2 array.

We tested different sizes of the phasing subsets as this was expected to have an effect on phasing accuracy. Tested values were 500, 1,000, 2,000, 5,000, and 10,000 individuals. For the Illumina 50Kv2 scenario we tested all combinations of core length and subset size. We only report subset size results for a fixed core length of 500 SNPs since the interaction between core length and subset size was minimal (data not shown). For the Illumina HD scenario we set the core length to 5,000 SNPs when testing subset size.

For evaluating the Heterogeneous Array scenarios we set the core length to 500 SNPs when the dataset consisted of only MD arrays since this value gave good performance in Homogeneous MD Array scenarios. Similarly, for datasets containing HD arrays we set core length to 5,000 SNPs. In both scenarios we set subset size to 5,000 SNPs.

AlphaPhase had several other parameters for which fixed default values were used. Specifically, we fixed the maximum number of surrogates used to ten and allowed 10% of the marker genotypes to disagree between pairs of surrogates. We also set the number of allele mismatches for clustering pairs of nearly identical library haplotypes to be zero.

When phasing multiple arrays we added an additional parameter to AlphaPhase. This parameter governs the minimum required number of matching alleles before two haplotypes can be identified as the same haplotype. If all SNPs were independent of each other, i.e., there was no linkage between them, we would expect the optimal value of this parameter to remain unchanged irrespective of SNP density. Our results (data not shown) suggest that the presence of linkage does not have a significant effect for the SNP densities considered here and that requiring a match of 200 alleles between two haplotypes is an appropriate value for this parameter. If SNP arrays with greater density are considered then the value of this parameter may need to be revised.

### Performance Testing

To test the performance of the new improvements to the LRP and HLI algorithms on large datasets we used the data from the Homogeneous Array scenarios for both the 100k and one million datasets. To test the scenario where parents are known and genotype information is available for them we evaluated phasing accuracy within each of the ten breeds individually using data from all generations. Similarly, to test the scenario where no parentage information is available we evaluated phasing accuracy for each of the ten generations individually (Figure 2). We report average results across either all ten families or all ten generations.

To test the speed and memory usage of AlphaPhase on large datasets we tested multiple combinations of number of generation and families from both the 1000k and the one million datasets using Homogeneous Array scenarios. To test the performance of the new improvements to the LRP and HLI algorithms on heterogeneous datasets we used the data from the Heterogeneous Array scenarios on the 100k dataset.

We report three phasing statistics – percentage of correctly phased alleles, percentage of unphased alleles, and percentage of incorrectly phased alleles. Unless explicitly stated otherwise, we report these statistics for heterozygous loci only. We also report on memory usage and runtimes. Runs were performed on computers with an Intel Xeon Processor E5-2630 v3 (2.4 GHz) and between 64 and 256GB of RAM.

## Results

### Long Range Phasing and Haplotype Library Imputation of Large Datasets

#### Core Length

To determine the accuracy of our new sub-setting method we first determined the optimal core length for each of the Illumina 50Kv2 and Illumina HD scenarios. Figure 3 and Tables S1-S2 show the accuracy on the Illumina 50Kv2 scenario for a variety of core lengths. Figure 3a shows the percentage of correctly phased heterozygous loci for the Illumina 50Kv2 array per family scenario. The percentage of correctly phased alleles increased as the core length increased, although the difference in accuracy between a core length of 500 SNPs (92.7%) and 1,000 SNPs (93.3%) was small. For the per generation scenario the percentage of correctly phased alleles peaked at a core length of 500 SNPs (92.3%) before dropping significantly for a core length of 1,000 SNPs (89.8%). The pattern for the number of incorrectly phased alleles for the Illumina 50Kv2 (Figure 3b) was less clear although there was a significant increase in the number of incorrectly phased alleles for a core length of 1,000 SNPs. Using a core length of 500 SNP the percentage of alleles incorrectly phased was 0.6% (per family scenario) and 0.9% (per generation scenario).

**Figure 3.**
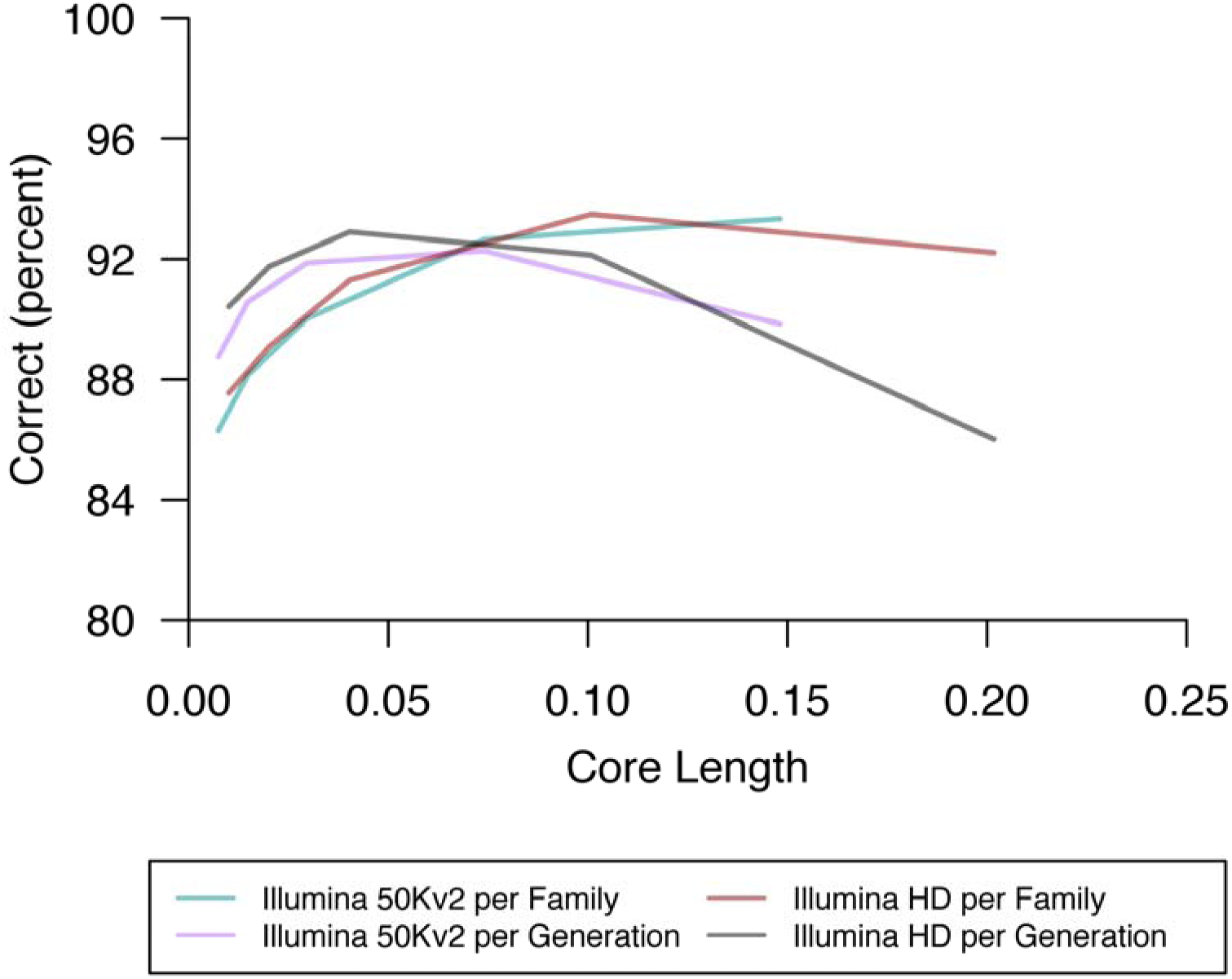
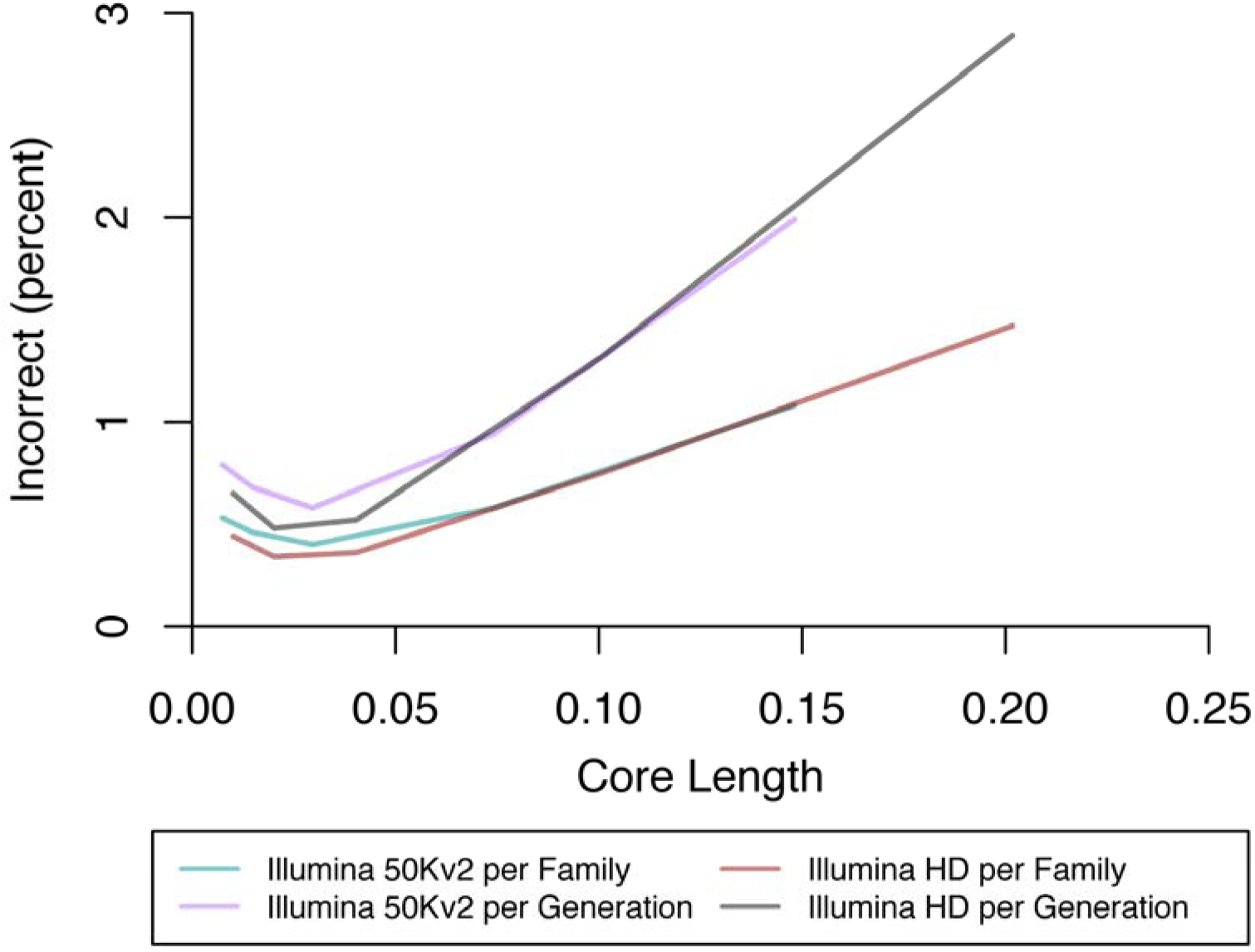
a) Percentage of correctly phased alleles at heterozygous loci for a range of core lengths. b) Percentage of incorrectly phased alleles at heterozygous loci for a range of core lengths. Core lengths are given as a proportion of the total chromosome length.

Figure 3 and Tables S3-S4 show that for the Illumina HD scenario the percentage of correctly phased alleles at heterozygous loci peaked at either a core length of 2,000 SNPs (per generation) or 5,000 SNPs (per family). For this scenario the number of incorrectly phased alleles was minimised at a core length of 1,000 SNPs (0.3% per family; 0.5% per generation), a shorter core length than that which maximised the number of correctly phased markers. Using a core length of 5,000 SNPs 93.5% (per family) or 92.1% (per generation) of alleles were phased correctly, while 0.8% (per family) or 1.3% (per generation) were phased incorrectly.

In all scenarios runtime was inversely proportional to core length (Tables S1-S4). We chose to study core lengths of 500 SNPs (for MD scenarios) and 5,000 SNPs (for HD scenarios) as a reasonable trade-off between accuracy and runtime. For these core lengths runtime was around three minutes for both arrays and for both the per generation and per family scenarios. Memory usage was 2.5GB for the Illumina 50Kv2 array and 5.3GB for the Illumina HD array (Tables S1-S4).

### Subset Size

Subset size can be expected to have a significant effect on the accuracy of phasing as it will directly influence the number of surrogates that are found. To test this we evaluated subset sizes of between 500 and 10,000 individuals (Figure 4, Tables S5-S8). For both the Illumina 50Kv2 and Illumina HD arrays accuracy increased as the subset size increased. For the Illumina 50Kv2 per family scenario the percentage of correctly phased alleles at heterozygous loci increased from 86.9% to 97.9% as subset size increased from 500 to 10,000 individuals. For the Illumina HD per family scenario it increased from 87.0% (500 individuals) to 97.8% (10,000 individuals). The results for phasing in the per generation scenarios were similar. The percentage of correctly phased alleles increased from 84.8% to 95.8% for the Illumina 50 Kv2 array and from 84.0% to 94.9% for the Illumina HD array as the subset size increased from 500 to 10,000 individuals.

**Figure 4.**
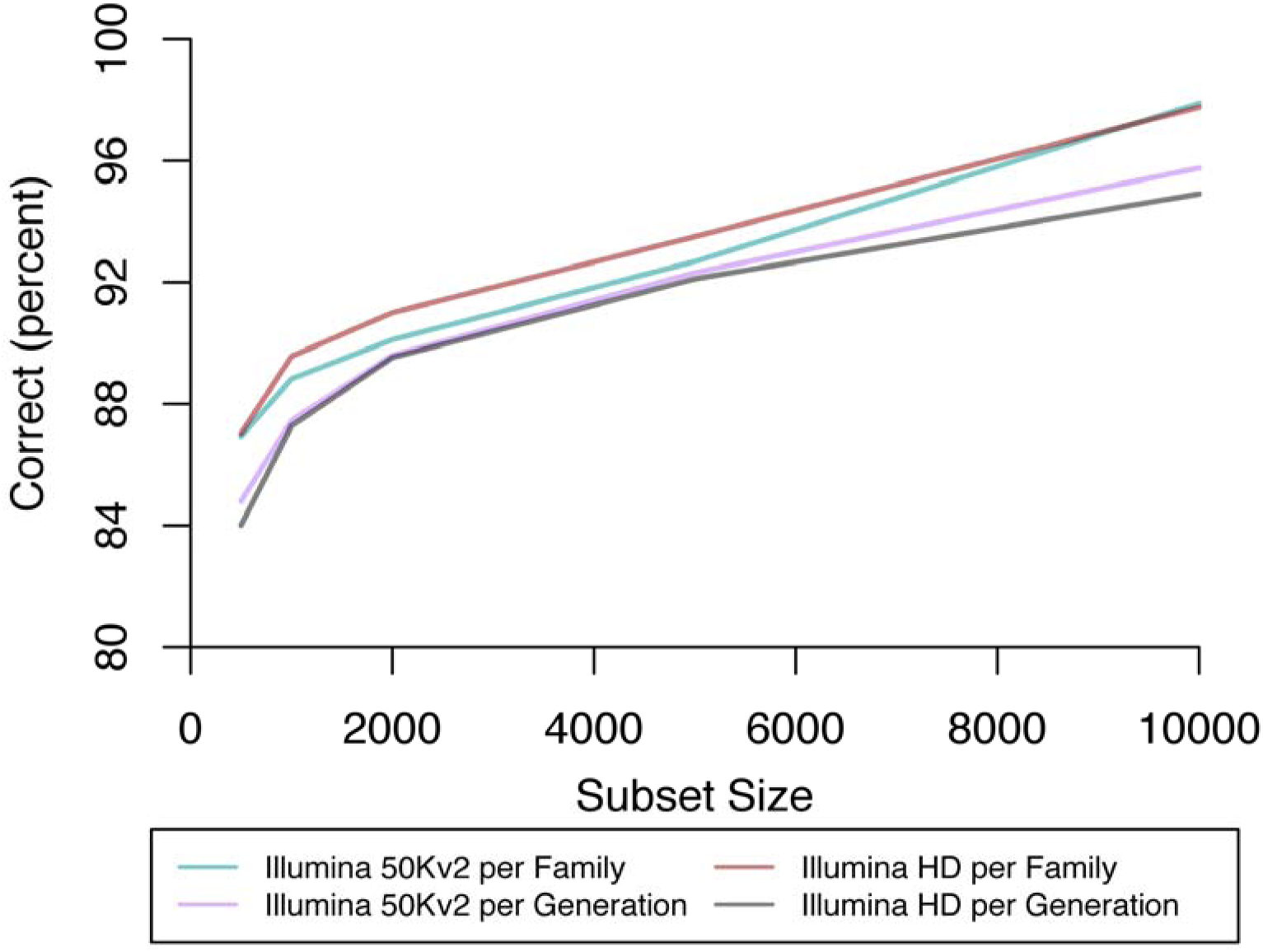
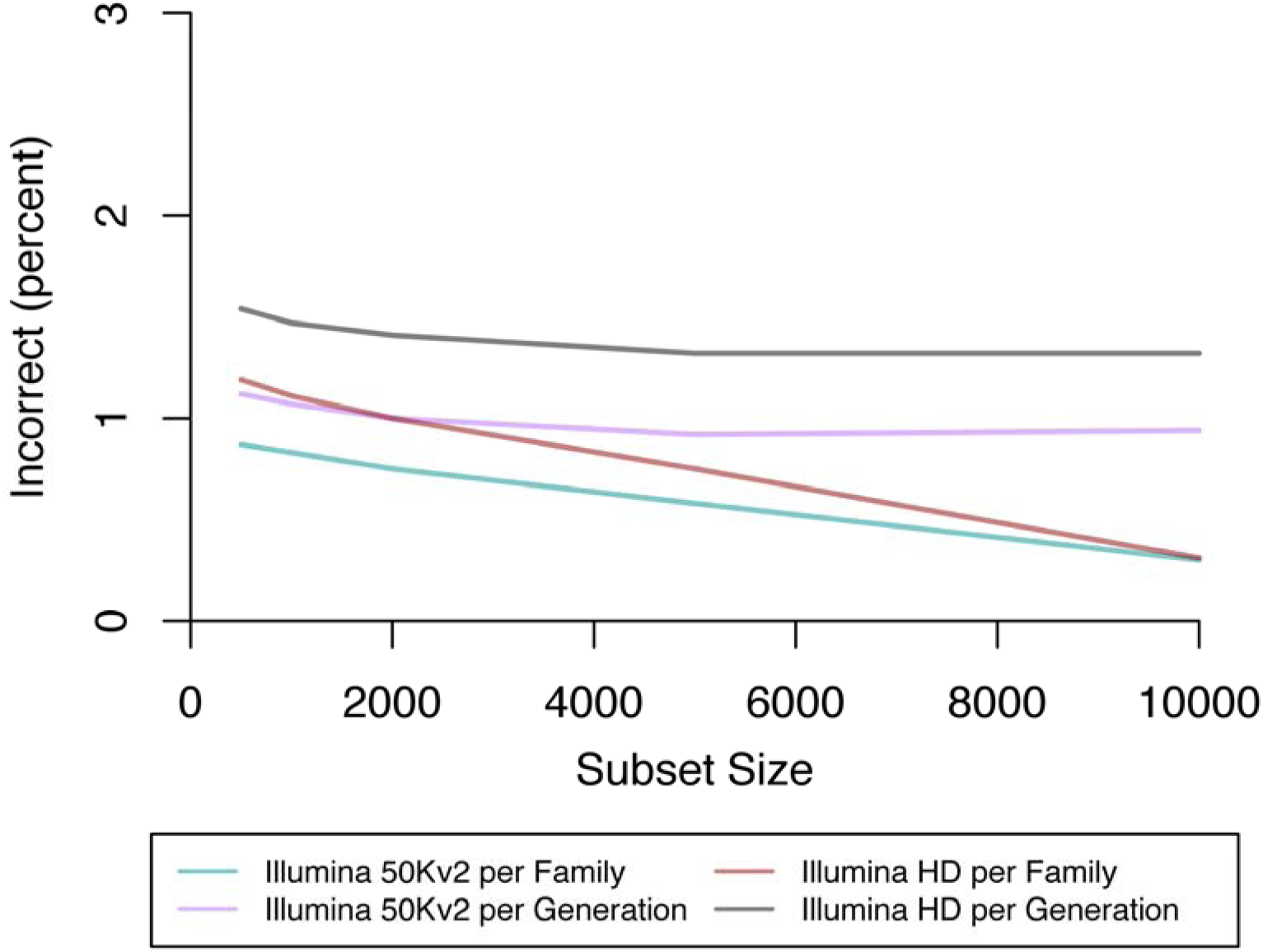
a) Percentage of correctly phased alleles at heterozygous loci for a range of subset sizes. b) Percentage of incorrectly phased alleles at heterozygous loci for a range of subset sizes.

In nearly all scenarios runtime was proportional to subset size (Tables S5-S8). The exception was for the Illumina 50Kv2 per family scenario in which runtime for a subset size of 10,000 individuals was noticeably shorter than runtime for a subset size of 5,000 individuals. This was likely due to nearly all animals having had both parents included in the subset when the subset size was 10,000, which can significantly decrease the time required to partition surrogates into maternal and paternal surrogates. Memory usage also increases as subset size grows. We chose to use a subset size of 5,000 for the remainder of this study as a reasonable trade-off between accuracy and runtime. With subsets of this size the percentage of correctly phased alleles at heterozygous loci was 92.7% (per family) or 92.3% (per generation) for the Illumina 50Kv2 scenario and 93.5% (per family) or 92.1% (per generation) for the Illumina HD scenarios. The percentage phased incorrectly was 0.1% (per family) or 0.9% (per generation) for the Illumina 50Kv2 scenarios and 0.8% (per family) or 1.3% (per generation) for the Illumina HD scenarios.

### Accuracy, Runtime, and Memory Usage on Different Dataset Sizes

To test the performance of our new improvements to the LRP and HLI algorithms on datasets of different sizes we created multiple different sized scenarios from the 100k and the one million datasets. Phasing accuracy was broadly comparable to the phasing accuracy observed when investigating optimal core length and subset size (Tables S9 and S10). Figure 5 shows that runtimes scaled approximately linearly with the number of individuals in a dataset. For the Illumina 50Kv2 dataset memory usage varied between 0.6GB for the smallest dataset of 1,000 individuals to 76GB for a dataset of one million individuals (Figure 6 and Table S19). Comparable figures for the Illumina HD dataset were 0.9GB and 325GB (Figure 6 and Table S10).

**Figure 5.**
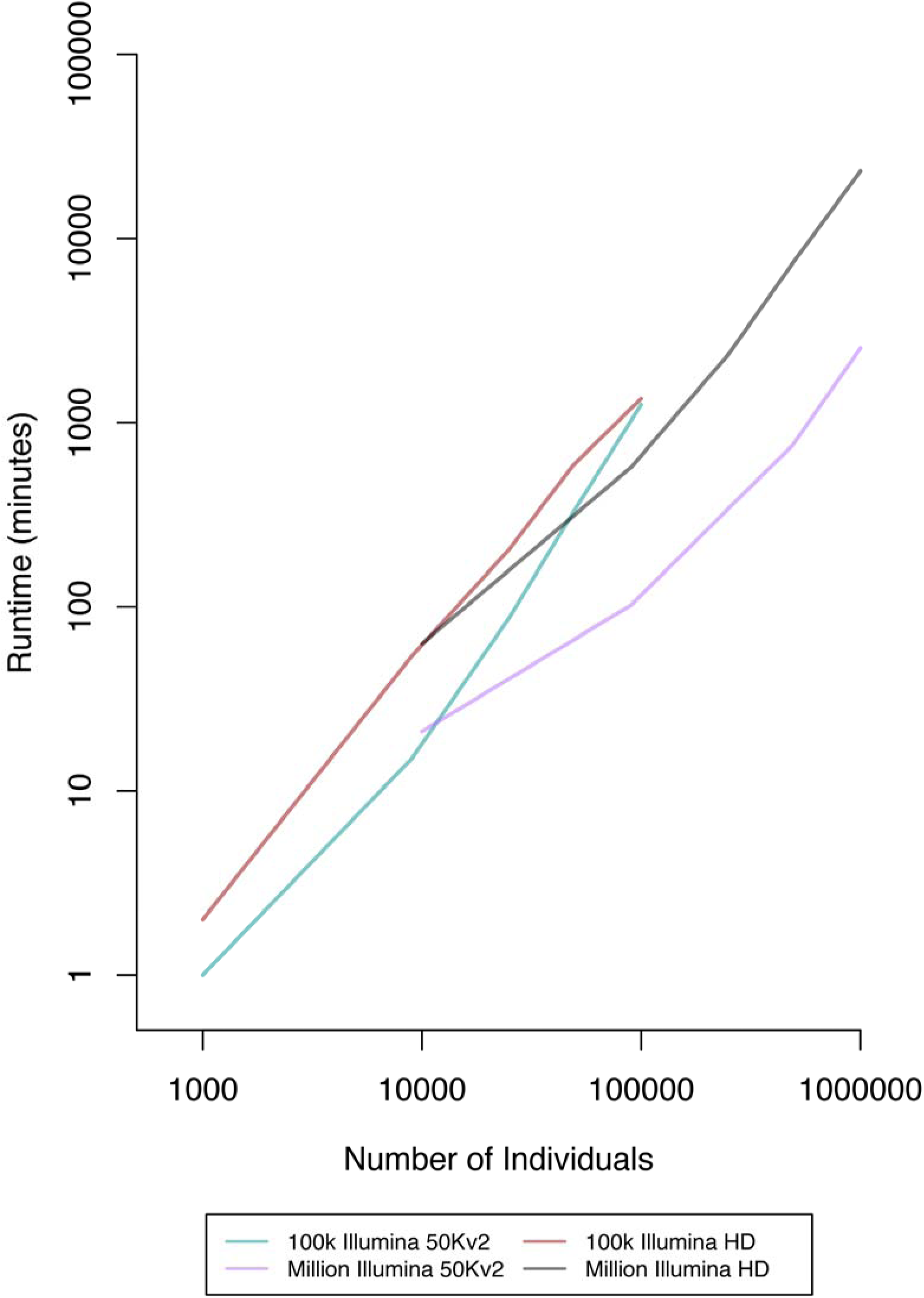
Runtime of AlphaPhase for a range of dataset sizes genotyped on two different SNP arrays.

**Figure 6.**
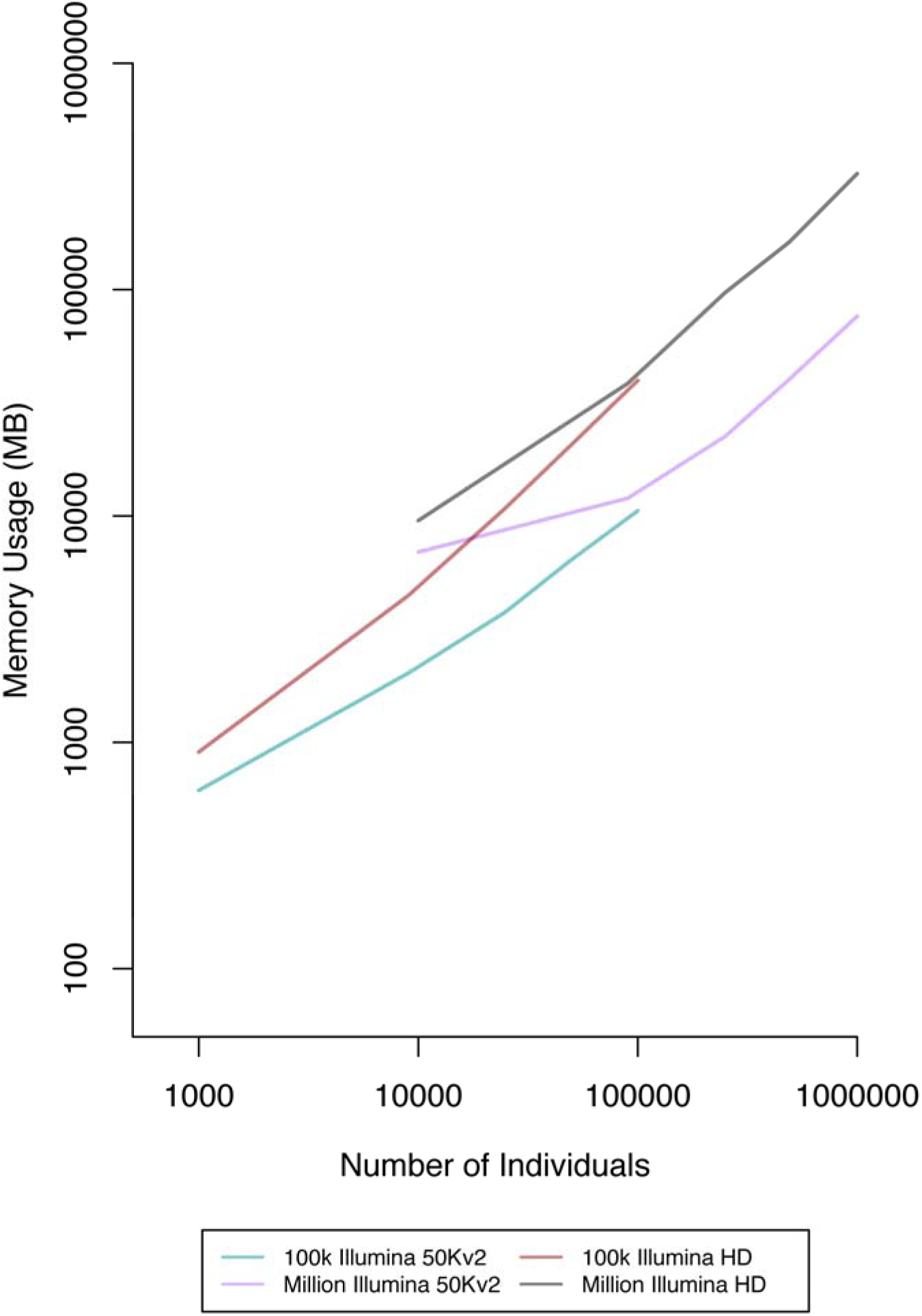
Memory usage of AlphaPhase for a range of dataset sizes genotyped on two different SNP arrays.

### Heterogeneous Datasets

Table 2 shows phasing accuracy, runtime and memory requirements for each of the five Heterogeneous Arrays per family scenarios. For the scenarios involving only MD arrays the percentage of alleles at heterozygous loci phased correctly was between 93.8% and 95.2% with between 1.9% and 2.8% phased incorrectly. For the two array scenarios (Two Illumina and Two Mixed) runtime was approximately two minutes. For the Three MD scenario runtime was approximately five minutes. For the two array scenarios memory usage was around 2.6GB whereas for the Three MD scenario memory usage was approximately 3GB.

**Table 2.** Different genotype scenarios per family results.

| Scenario | All |  |  | Heterozygous |  |  | Time<br>(minutes) | Memory<br>(MB) |
| --- | --- | --- | --- | --- | --- | --- | --- | --- |
|  | Correct | Unphased | Incorrect | Correct | Unphased | Incorrect |  |  |
| Illumina<br>50Kv2 | 98.30 | 1.59 | 0.11 | 92.65 | 6.77 | 0.58 | 143 | 2,585 |
| Illumina HD | 98.40 | 1.46 | 0.14 | 93.48 | 5.77 | 0.75 | 154 | 5,279 |
| Two Illumina | 94.12 | 3.18 | 2.70 | 93.83 | 4.25 | 1.92 | 126 | 2,570 |
| Two Mixed | 96.67 | 2.77 | 0.56 | 94.41 | 3.26 | 2.33 | 153 | 2,609 |
| Three MD | 96.66 | 2.41 | 0.93 | 95.15 | 2.05 | 2.79 | 322 | 3,034 |
| Mixed MD/HD | 93.64 | 5.12 | 1.23 | 93.41 | 4.38 | 2.20 | 342 | 5,311 |
| Ten Array | 95.12 | 4.05 | 0.83 | 95.37 | 3.66 | 0.97 | 562 | 7,671 |

We also tested a mixture of one MD array and one HD array with nine individuals genotyped on the MD array for every individual genotyped on the HD array (Mixed MD/HD). As expected the percentage of correctly phased alleles at heterozygous loci was lower than in other scenarios, but was still 93.4%. Runtime was around six hours and memory usage was 5.3GB. For the Ten Array scenario the percentage of correctly phased alleles was slightly higher at 95.4%, although memory usage was also higher at 7.6GB.

Table 3 shows results for the per generation scenarios. Results were broadly comparable to the per family results. Across the scenarios containing only MD arrays the percentage of correctly phased alleles at heterozygous loci was between 93.1% and 95.6% with between 3.1% and 3.6% incorrectly phased. Runtime was similar to the per family scenarios taking around three minutes for the two array scenarios and two minutes for the Three MD scenarios. Memory usage was the same as that of the per family scenarios.

**Table 3.**
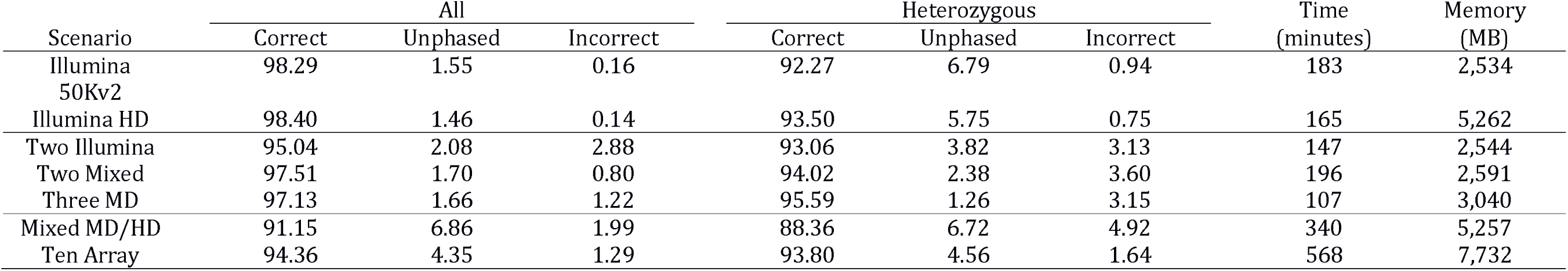
Different genotype scenarios per generation results.

The Mixed HD/MD per generation scenario showed a lower percentage of correctly phased alleles at heterozygous loci (88.4%) compared to the per family scenario (93.4%). Runtime and memory requirements were similar. The Ten Array per generation scenario also had lower accuracy than the per family scenario with 93.8% of alleles correctly phased compared with 95.4% in the per family scenario. Runtime and memory usage were similar for the per family and the per generation results.

## Discussion

In this paper we introduced improvements to the LRP and HLI algorithms of AlphaPhase [16] to enable phasing of very large and heterogeneous datasets in which individuals have been genotyped on differing sets of markers. We tested the revised algorithms’ performance on a range of simulated datasets and show that AlphaPhase can be used to accurately phase datasets that contain up to one million individuals and that have been genotyped with multiple different SNP arrays. In what follows we discuss the effect of: (i) core length and (ii) subset size on phasing accuracy and computational runtime and memory use; and (iii) the impact of these improvements on the phasing of large and heterogeneous datasets.

### Effect of core length on phasing performance

Both LRP and HLI break the genome into smaller sections of consecutive SNPs called cores. Each of these cores is then phased independently of each other. Core length, defined as the number of SNPs in each core, has previously been shown to have a significant effect on accuracy [16]. In light of our new improvements to the LRP and HLI algorithms and the availability of much denser SNP arrays we further investigated the impact of this parameter.

As expected we found that core length has a significant effect on phasing accuracy. Short cores showed similar levels of phasing accuracy. Accuracy started to deteriorate notably as cores get longer. We also found that phasing accuracy is a function of the length of the core as a proportion of the length of the chromosome, rather than the number of SNPs it contains, and that the Illumina 50Kv2 and Illumina HD arrays show a very similar pattern of results when core length is expressed as a proportion of chromosome length. This is to be expected, as phasing accuracy is likely to be highly affected by the presence of recombinations within a core. The latter can be reliably measured by the relative size of a core versus the whole chromosome, and less so by the number of SNP array markers in a core. The reduction in accuracy observed as the core length increased is likely due to the increased chance of a core containing a recent recombination. This would reduce the number of surrogates and thus, reduce the information available for phasing.

Both runtime and memory usage were significantly affected by core length with runtime being approximately proportional to the number of cores and therefore, inversely proportional to core length. Memory usage also increased as the number of cores increased although the effect was less pronounced than for runtime. For these reasons we recommend the use of the longest possible cores that do not result in an unacceptable drop in accuracy. For the bovine arrays considered in this paper, our results suggest a core length of 500 when only MD arrays are used and a core length of 5,000 when HD arrays have also been used.

### Effect of subset size on phasing performance

Our new improvements of the LRP algorithm partition a large dataset into subsets and then performing LRP on each of the subsets. We introduced a new parameter to the LRP algorithm that controls the size of these subsets and our results show that this parameter can have a significant effect on phasing accuracy. A subset size of 10,000 gave both the highest percentage of correctly phased alleles and the lowest percentage of incorrectly phased ones. As the subset size decreased the proportion of correctly phased alleles decreased and the proportion of incorrectly phased alleles loci increased. This decrease in phasing performance was very likely due to the reduction in the number of surrogates that would be expected in a smaller subset which, in turn, led to less information with which to accurately phase.

The increase in phasing accuracy that resulted from increasing subset size had a significant cost in terms of runtime. We found an approximately linear relationship between the size of the subset and runtime with the used runtime parameters. Consequently, there was a trade-off between runtime and accuracy. As the size of subsets got larger the number of incorrectly phased alleles appeared to begin to plateau for a subset size of 5,000 or greater and runtime started to increase significantly. For the datasets we tested the subset size for which this value occurred seemed to be largely invariant to total dataset size and so we suggest that a subset size of 5,000 is an appropriate compromise. The optimal size for datasets with a different structure, such as a human populations in which individuals are likely to be less related than the ones considered here, warrants further investigation.

Phasing of large datasets is likely to be computationally expensive for any phasing techniques due to the large number of individuals involved. Our results suggest that there is sufficient information in small subsets from a larger dataset to allow a significant number of alleles to be phased accurately. This suggests that for other phasing methods, such as those based on probabilistic models, it could also be beneficial to break the phasing of large datasets into subsets before merging the results.

### Ability of AlphaPhase to phase large heterogeneous datasets

Our results show that it is viable to run heuristic phasing on very large datasets, such as those now available for humans [10] or cattle [4,11,12]. AlphaPhase took two days and 76GB of memory to phase one million animals genotyped on a simulated Illumina 50Kv2 array. To phase one million animals genotyped on a simulated Illumina HD array took 14 days and 325GB of memory.

The ability to phase datasets genotyped using multiple different arrays is important as datasets are increasingly likely to consist of individuals genotyped using different arrays due to the increase in the number of available arrays for commonly genotyped species. Results from the analysis of the Heterogenous Arrays scenarios show that similar phasing accuracy can be achieved for heterogenous datasets, consisting of individuals genotyped on multiple MD arrays, as can be achieved for homogeneous datasets. In general, accuracy was slightly worse than for the single array scenarios that were tested, although in many scenarios the Heterogeneous Arrays phased slightly more alleles correctly. However, this increase in percentage of correctly phased alleles came at the cost of phasing more alleles incorrectly as well.

AlphaPhase can now also accurately phase individuals genotyped on a mixture of MD and HD SNP arrays. The phasing of such datasets is likely to become increasingly common as it is desirable to continue to use the data already collected using MD arrays even as the use of HD arrays grows. Although the phasing accuracy for heterogeneous datasets was often lower than when individuals were genotyped on a single SNP array, the percentage of correctly phased alleles was still over 93% in all scenarios tested other than the Mixed MD/HD per generation scenario. In this scenario many individuals had high amounts of missing data due to them being genotyped on the MD rather than HD array and so we would expect phasing to be more difficult.

## Conclusions

We have modified the LRP and HLI algorithms to allow phasing of large heterogeneous datasets. These modifications are implemented in AlphaPhase version 1.3.7 (available from http://alphagenes.roslin.ed.ac.uk/) and allow the accurate phasing of millions of individuals genotyped on multiple SNP arrays.

## Declarations

### Ethics approval and consent to participate

Not applicable

### Consent for publication

Not applicable

### Availability of data and materials

An implementation of our algorithm, in the software package AlphaPhase is available from the authors’ website, https://alphagenes.roslin.ed.ac.uk/wp/software/alphaphase/, and is free for academic use.

### Authors’ contributions

JMH and DM designed the updates to the algorithm and this study. DM and DW implemented the updates. JJ and GG provided the simulation strategy. DM conducted the study. GG provided critical comments throughout the study. All authors read and approved the final manuscript.

## Acknowledgements

This work has made use of the resources provided by the Edinburgh Compute and Data Facility (ECDF) (http://www.ecdf.ed.ac.uk).

## Supplementary tables

**Table S1:** Illumina 50Kv2 per family results for a range of core lengths.

| Core Length | All Loci |  |  | Heterozygous Loci |  |  | Time | Memory |
| --- | --- | --- | --- | --- | --- | --- | --- | --- |
|  | Correct | Unphased | Incorrect | Correct | Unphased | Incorrect | (minutes) | (MB) |
| 50 | 97.37 | 2.53 | 0.10 | 86.30 | 13.16 | 0.53 | 3,247 | 5,930 |
| 100 | 97.66 | 2.25 | 0.09 | 88.14 | 11.40 | 0.46 | 1,392 | 4,286 |
| 200 | 97.93 | 2.00 | 0.07 | 90.02 | 9.58 | 0.40 | 561 | 3,271 |
| 500 | 98.30 | 1.59 | 0.11 | 92.65 | 6.77 | 0.58 | 143 | 2,585 |
| 1,000 | 98.39 | 1.40 | 0.21 | 93.33 | 5.60 | 1.08 | 39 | 2,060 |

**Table S2:** Illumina 50Kv2 per generation results for a range of core lengths.

| Core Length | All Loci |  |  | Heterozygous Loci |  |  | Time | Memory |
| --- | --- | --- | --- | --- | --- | --- | --- | --- |
|  | Correct | Unphased | Incorrect | Correct | Unphased | Incorrect | (minutes) | (MB) |
| 50 | 97.92 | 1.93 | 0.15 | 88.75 | 10.46 | 0.79 | 4,017 | 5,946 |
| 100 | 98.18 | 1.69 | 0.13 | 90.57 | 8.76 | 0.68 | 1,729 | 4,263 |
| 200 | 98.34 | 1.56 | 0.10 | 91.87 | 7.55 | 0.58 | 706 | 3,233 |
| 500 | 98.29 | 1.55 | 0.16 | 92.27 | 6.79 | 0.94 | 183 | 2,534 |
| 1,000 | 97.82 | 1.84 | 0.34 | 89.84 | 8.17 | 1.99 | 46 | 2,047 |

**Table S3:**
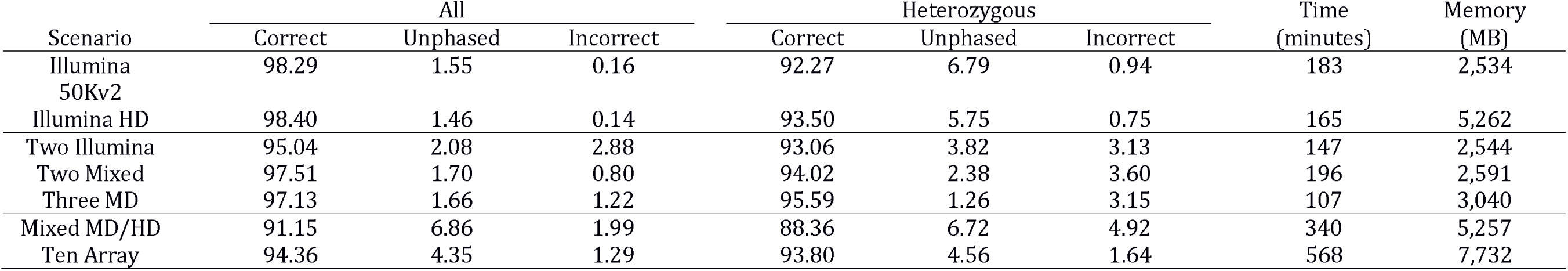
Illumina H D per family results for a range of core lengths.

**Table S4:** Illumina HD per generation results for a range of core lengths.

| Core Length | All Loci |  |  | Heterozygous Loci |  |  | Time | Memory |
| --- | --- | --- | --- | --- | --- | --- | --- | --- |
|  | Correct | Unphased | Incorrect | Correct | Unphased | Incorrect | (minutes) | (MB) |
| 500 | 98.16 | 1.71 | 0.12 | 90.42 | 8.93 | 0.65 | 3,138 | 7,783 |
| 1,000 | 98.32 | 1.59 | 0.09 | 91.75 | 7.77 | 0.48 | 1,351 | 6,683 |
| 2,000 | 98.43 | 1.48 | 0.09 | 92.90 | 6.58 | 0.52 | 583 | 5,990 |
| 5,000 | 98.19 | 1.59 | 0.23 | 92.13 | 6.55 | 1.32 | 183 | 5,255 |
| 10,000 | 97.19 | 2.31 | 0.50 | 86.01 | 11.10 | 2.89 | 79 | 4,877 |

**Table S5:** Illumina 50Kv2 per family results for a range of subset sizes.

| Subset Size | All Loci |  |  | Heterozygous Loci |  |  | Time | Memory |
| --- | --- | --- | --- | --- | --- | --- | --- | --- |
|  | Correct | Unphased | Incorrect | Correct | Unphased | Incorrect | (minutes) | (MB) |
| 500 | 96.98 | 2.86 | 0.16 | 86.93 | 12.20 | 0.16 | 4 | 1,246 |
| 1,000 | 97.39 | 2.45 | 0.16 | 88.82 | 10.34 | 0.16 | 8 | 1,360 |
| 2,000 | 97.69 | 2.17 | 0.14 | 90.11 | 9.13 | 0.14 | 23 | 2,153 |
| 5,000 | 98.31 | 1.58 | 0.11 | 92.68 | 6.75 | 0.11 | 138 | 2,579 |
| 10,000 | 99.60 | 0.34 | 0.06 | 97.89 | 1.81 | 0.06 | 56 | 3,277 |

**Table S6:** Illumina 50Kv2 per generation results for a range of subset sizes.

| Subset Size | All Loci |  |  | Heterozygous Loci |  |  | Time | Memory |
| --- | --- | --- | --- | --- | --- | --- | --- | --- |
|  | Correct | Unphased | Incorrect | Correct | Unphased | Incorrect | (minutes) | (MB) |
| 500 | 96.40 | 3.40 | 0.20 | 84.80 | 14.07 | 1.12 | 4 | 1,248 |
| 1,000 | 97.00 | 2.81 | 0.19 | 87.45 | 11.48 | 1.07 | 8 | 1,367 |
| 2,000 | 97.55 | 2.27 | 0.17 | 89.60 | 9.40 | 1.00 | 24 | 1,955 |
| 5,000 | 98.30 | 1.54 | 0.16 | 92.31 | 6.77 | 0.92 | 168 | 2,530 |
| 10,000 | 99.28 | 0.56 | 0.16 | 95.76 | 3.30 | 0.94 | 703 | 3,788 |

**Table S7:** Illumina HD per family results for a range of subset sizes.

| Subset Size | All Loci |  |  | Heterozygous Loci |  |  | Time | Memory |
| --- | --- | --- | --- | --- | --- | --- | --- | --- |
|  | Correct | Unphased | Incorrect | Correct | Unphased | Incorrect | (minutes) | (MB) |
| 500 | 96.94 | 2.83 | 0.22 | 87.03 | 11.78 | 1.19 | 20 | 4,146 |
| 1,000 | 97.45 | 2.34 | 0.21 | 89.56 | 9.33 | 1.11 | 30 | 4,243 |
| 2,000 | 97.76 | 2.05 | 0.19 | 91.00 | 8.00 | 1.00 | 49 | 4,854 |
| 5,000 | 98.40 | 1.46 | 0.14 | 93.50 | 5.75 | 0.75 | 165 | 5,262 |
| 10,000 | 99.57 | 0.37 | 0.06 | 97.76 | 1.94 | 0.31 | 247 | 6,109 |

**Table S8:** Illumina HD per generation results for a range of subset sizes.

| Subset Size | All Loci |  |  | Heterozygous Loci |  |  | Time | Memory |
| --- | --- | --- | --- | --- | --- | --- | --- | --- |
|  | Correct | Unphased | Incorrect | Correct | Unphased | Incorrect | (minutes) | (MB) |
| 500 | 96.20 | 3.54 | 0.27 | 84.01 | 14.45 | 1.54 | 19 | 4,172 |
| 1,000 | 96.86 | 2.89 | 0.25 | 87.27 | 11.26 | 1.47 | 30 | 4,280 |
| 2,000 | 97.43 | 2.33 | 0.24 | 89.52 | 9.07 | 1.41 | 48 | 4,759 |
| 5,000 | 98.18 | 1.59 | 0.23 | 92.11 | 6.57 | 1.32 | 163 | 5,261 |
| 10,000 | 99.12 | 0.65 | 0.22 | 94.90 | 3.78 | 1.32 | 533 | 6,586 |

**Table S9:** Illumina 50Kv2 results for scenarios of different sizes.

| Dataset | Families × | Number of | All Loci |  |  | Heterozygous Loci |  |  | Time | Memory |
| --- | --- | --- | --- | --- | --- | --- | --- | --- | --- | --- |
|  | Generations | Individuals | Correct | Unphased | Incorrect | Correct | Unphased | Incorrect | (minutes) | (MB) |
| 100k | 1 × 1 | 1,000 | 99.62 | 0.30 | 0.07 | 98.39 | 1.29 | 0.31 | 1 | 613 |
| 100k | 3 × 3 | 9,000 | 98.08 | 1.86 | 0.06 | 93.19 | 6.55 | 0.26 | 15 | 2,022 |
| 100k | 5 × 5 | 25,000 | 98.01 | 1.89 | 0.10 | 92.59 | 6.96 | 0.45 | 87 | 3,770 |
| 100k | 7 × 7 | 49,000 | 97.98 | 1.88 | 0.14 | 91.82 | 7.50 | 0.67 | 326 | 6,326 |
| 100k | 10 × 10 | 100,000 | 97.81 | 2.02 | 0.17 | 90.25 | 8.83 | 0.92 | 1,253 | 10,542 |
| One million | 1 × 1 | 10,000 | 98.59 | 1.35 | 0.06 | 95.48 | 4.27 | 0.25 | 21 | 6,927 |
| One million | 3 × 3 | 90,000 | 96.98 | 2.94 | 0.07 | 89.73 | 9.95 | 0.32 | 102 | 11,972 |
| One million | 5 × 5 | 250,000 | 97.62 | 2.27 | 0.11 | 91.40 | 8.13 | 0.47 | 341 | 22,455 |
| One million | 7 × 7 | 490,000 | 97.70 | 2.17 | 0.13 | 91.30 | 8.10 | 0.60 | 751 | 40,112 |
| One million | 10 × 10 | 1,000,000 | 97.60 | 2.22 | 0.18 | 90.35 | 8.84 | 0.81 | 2,534 | 76,305 |

**Table S10:**
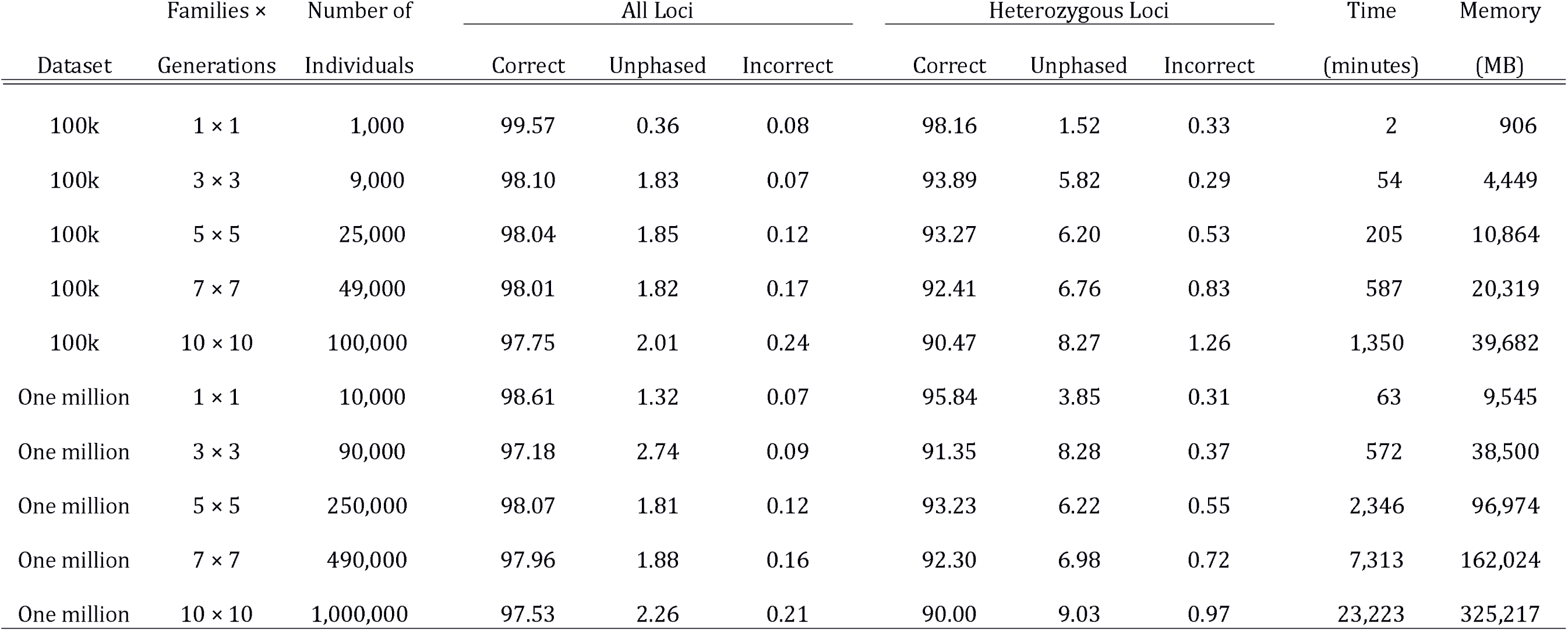
Illumina HD results for scenarios of different sizes.

| Dataset | Families × | Number of | All Loci |  |  | Heterozygous Loci |  |  | Time | Memory |
| --- | --- | --- | --- | --- | --- | --- | --- | --- | --- | --- |
|  | Generations | Individuals | Correct | Unphased | Incorrect | Correct | Unphased | Incorrect | (minutes) | (MB) |
| 100k | 1 × 1 | 1,000 | 99.57 | 0.36 | 0.08 | 98.16 | 1.52 | 0.33 | 2 | 906 |
| 100k | 3 × 3 | 9,000 | 98.10 | 1.83 | 0.07 | 93.89 | 5.82 | 0.29 | 54 | 4,449 |
| 100k | 5 × 5 | 25,000 | 98.04 | 1.85 | 0.12 | 93.27 | 6.20 | 0.53 | 205 | 10,864 |
| 100k | 7 × 7 | 49,000 | 98.01 | 1.82 | 0.17 | 92.41 | 6.76 | 0.83 | 587 | 20,319 |
| 100k | 10 × 10 | 100,000 | 97.75 | 2.01 | 0.24 | 90.47 | 8.27 | 1.26 | 1,350 | 39,682 |
| One million | 1 × 1 | 10,000 | 98.61 | 1.32 | 0.07 | 95.84 | 3.85 | 0.31 | 63 | 9,545 |
| One million | 3 × 3 | 90,000 | 97.18 | 2.74 | 0.09 | 91.35 | 8.28 | 0.37 | 572 | 38,500 |
| One million | 5 × 5 | 250,000 | 98.07 | 1.81 | 0.12 | 93.23 | 6.22 | 0.55 | 2,346 | 96,974 |
| One million | 7 × 7 | 490,000 | 97.96 | 1.88 | 0.16 | 92.30 | 6.98 | 0.72 | 7,313 | 162,024 |
| One million | 10 × 10 | 1,000,000 | 97.53 | 2.26 | 0.21 | 90.00 | 9.03 | 0.97 | 23,223 | 325,217 |

